# Systemic administration of anti-CD20 indirectly reduces B cells in the inflamed meninges in a chronic model of central nervous system autoimmunity

**DOI:** 10.1101/2021.01.22.427667

**Authors:** Yodit Tesfagiorgis, Heather C Craig, Kate A Parham, Steven M Kerfoot

**Affiliations:** Department of Microbiology and Immunology University of Western Ontario, London, Ontario, Canada

**Keywords:** B cell, T cell, autoimmunity, EAE, MOG, Multiple Sclerosis, anti-CD20

## Abstract

Anti-CD20 B cell depleting therapies have demonstrated that B cells are important drivers of disease progress in Multiple Sclerosis, although the pathogenic mechanisms are not well understood. A population of B cells accumulates in the inflamed meninges in MS and also some chronic animal models of disease, typically adjacent to demyelinating lesions. The role of these meningeal B cells in disease is not known, nor is their susceptibility to anti-CD20 therapy. Here, we administered anti-CD20 to 2D2 IgH^MOG^ spontaneous experimental autoimmune encephalomyelitis mice in the chronic phase of disease, after the establishment of meningeal B cell clusters. Compared to the circulation, lymph nodes, and spleen, B cell depletion from the CNS was delayed and not evident until 7d post administration of anti-CD20. Further, we did not find evidence that anti-CD20 accessed meningeal B cells directly, but rather that depletion was indirect and the result of ongoing turnover of the meningeal population and elimination of the peripheral pool from which it is sustained. The reduction of B cell numbers in the CNS coincided with less demyelination of the spinal cord white matter and also, surprisingly, an increase in the number of T cells recruited to the meninges but not parenchyma.

## Introduction

Anti-CD20 B cell depleting therapies, which includes rituximab, ocrelizumab, and ofatumumab, are effective in treating patients with relapsing(1–3) and progressive forms of multiple sclerosis (MS) (4–6). In clinical trials anti-CD20 was shown to reduce the number of B cells in the cerebral spinal fluid (CSF) (6–9) and peripheral blood (3, 10) and also results in a reduction of important measures of disease activity including a reduction in new and existing brain lesions, clinical relapses, disability progression (1–5, 10–12) and, importantly, T cell accumulation in the CSF (9) and peripheral blood (8). The B cell lineage is best known for their production of antibodies that target immune effector mechanisms to specific antigen targets. However, because anti-CD20 therapies do not target antibody producing plasma cells, nor do they reduce antibody levels within the therapeutic time frame (2, 8, 9), it is not clear how B cells are contributing to disease pathogenesis.

B cells accumulate in the central nervous system (CNS) of MS patients, where they often form clusters within the meninges directly adjacent to demyelinating lesions (13–16). Importantly, these clusters have been shown to correlate to early onset of disease and severe cortical pathology (14, 15). We and others have shown that this phenomenon is recapitulated in some animal models of anti-myelin autoimmunity (experimental autoimmune encephalomyelitis - EAE) (17–21). These clusters can become organized with separate T cell zones and B cell follicles reminiscent of lymphoid tissue (13, 22), but more frequently they are disorganized mixtures of B and T cells (23, 24). Because of their location within the inflamed CNS, it has been hypothesized that these lymphoid clusters represent an environment where the autoimmune response is propagated locally. Nevertheless, this has not yet been demonstrated and the pathogenic contributions to local disease remain unknown. Further, it is not yet known if B cells in the meningeal clusters are susceptible to depletion with systemically administered anti-CD20 antibodies used to successfully treat MS.

B cells play multiple roles in MS and EAE, complicating the analysis of the therapeutic benefits of B cell depletion. Work by ourselves and others suggests that, depending on the stage of disease, both myelin-specific and non-specific B cells contribute different mechanisms to disease. For instance, in a peptide induced model of EAE, depletion of all B cells, including non-specific cells, prior to disease induction results in more severe disease (25, 26). This is attributed to IL-10 production by B cells, demonstrating that at least some B cells can play a regulatory role. In contrast, anti-myelin B cells contribute to disease initiation in models that incorporate B cells in the autoimmune response. This includes 2D2 IgH^MOG^ spontaneous EAE (sEAE) (27, 28) and models induced by immunization with larger protein myelin antigens (26, 29, 30). Indeed, EAE induced in mice by immunization with human myelin oligodendrocyte glycoprotein (MOG) is completely dependent on B cells through the production of anti-MOG antibodies (31). Nevertheless, we have shown that the B cell anti-myelin response is short-lived (32) and that anti-myelin B cells are not found in meningeal clusters in the inflamed spinal cord in either sEAE or MOG-protein induced EAE (33). Therefore, the role of anti-myelin-specific B cells may be confined to the initiation of the disease, and thus the contribution of B cells to the chronic phase of disease is likely from some other, non-myelin specific population and may include those that accumulate in the CNS.

Here, we set out to determine if B cells in meningeal clusters are susceptible to depletion by systemic anti-mCD20 treatment. We specifically investigated B cell depletion following treatment in the chronic phase of disease, after the establishment of B and T cell inflammation in the CNS, in order to isolate the pathogenic contribution of these cells from any role B cells play in disease initiation. Further, considering the association tween meningeal B cells and severe disease, we wanted to determine if a reduction in these cells did correspond to a reduction in pathology. To address these questions, we employed the 2D2 IgH^MOG^ sEAE model which we have shown results in a chronic disease course with consistent and robust, ongoing inflammation in the spinal cord that includes B cell accumulation in the meninges adjacent to demyelinating lesions (20). We found that systemic administration of anti-mCD20 does not target B cells in meningeal clusters directly, but does reduce their numbers over time, likely due to the elimination of peripheral B cells and the ongoing turnover of the meningeal population. Further, we found that the reduction in B cells in meningeal clusters corresponds to reduced demyelination in the adjacent lesion. Finally, and unexpectedly, B cell depletion also corresponded to an increase in T cells specifically in the meninges, but not white matter.

## Materials and Methods

### Mice

Wild-type C57BL/6 and 2D2 TCR transgenic (211) were purchased from the Jackson Laboratory. Mice expressing fluorescent dsRed (6051; Tg(CAG-DsRed∗MST)1Nagy/J) under control of the β-Actin promoter within all nucleated cells were obtained from the Jackson Laboratory. IgH^MOG^ MOG-specific BCR knockin mice (212) were received as a gift from Dr. H. Wekerle. All mice were housed under specific pathogen-free conditions at the West Valley Barrier Facility at Western University Canada. Animal protocols were approved by the Western University Animal Use Subcommittee.

### Spontaneous 2D2 IgH^MOG^ EAE model

IgH^MOG+/+^ mice were crossed with 2D2^+/-^ mice. Approximately 80% of IgH^MOG+/-^ 2D2^+/-^ double mutant offspring spontaneously developed signs of EAE (sEAE) at 31 – 42d of age. Only mice that developed signs of disease and met the conditions of chronic disease were included in experiments. As both sexes develop chronic disease, both male and female mice were used in experiments. Clinical disease was monitored daily with a modified 0-20 clinical scoring system to evaluate tail paralysis, weakness and paralysis for each individual limb, and righting reflex. Scores were determined as follows: Tails were scored as: 0, asymptomatic; 2, partial tail paralysis; 4, complete tail paralysis. Each hind limb was scored as: 0, asymptomatic; 1, hind limb weakness with a wobbly gate; 2, weight bearing but knuckling; 3, not weight bearing; 4, complete hind limb paralysis. Each forelimb was scored as: 0, asymptomatic; 1, weak grasp yet weight bearing; 2, not weight bearing; 3, complete forelimb paralysis. Finally, the righting reflex was assessed as: 0, asymptomatic; 1, delayed righting reflex; 2, no righting reflex.

### Antibodies for flow cytometry and histology

The following Abs were purchased from BD Bioscience: anti-CD4-v450 (RM4-5), anti-CD45R-A647 (RA3-6B2), anti-CD138-BV421 (281-2), CD19-BV711 (1D3) and mIgG2a-biotin (R19-15). The following Abs were purchased from BioLegend: anti-CD4-A647 (RM4-5), anti-CD3-A488 (17A2), CD45R-APC-Cy7 (RA3-6B2), anti-IgKappa-biotin (RMK-12), anti-Ly6G-A647 (1A8), anti-rabbit-DyLight555 (Poly4064), anti-CD19-A488 (6D5) and anti-GFAP-A488 (2E1.E9). Streptavidin-APC, Streptavidin-eF570 and anti-CD4-PECy5 (RM4-5) was purchased from eBioscience. Anti-Myelin Basic Protein (MBP)-rabbit polyclonal antibody and anti-Ki67 (SP6) unconjugated was purchased from Abcam.

### Anti-mCD20 B cell depletion

Anti-mouse-CD20 (5D2), the murine surrogate of rituximab was received as a generous gift from Genentech, South San Francisco, USA. 150μg of the drug was administered i.v. (unless otherwise stated) during the chronic phase of disease, ~2 weeks post disease onset. Anti-IgG2a (MG2a-53) acquired from BioLegend was used as an isotype control and injected at the same concentration i.v. Mice were monitored for EAE disease severity for either 2- or 7-days following treatment before collecting the blood, lymph node, spleen and spinal cord for flow cytometry or histology.

### Flow cytometry

The blood, lymph nodes (inguinal, axillary, and cervical), spleen and the spinal cord were harvested from mice for flow cytometry analysis as previously described (18, 33). Briefly, blood was isolated through a cardiac puncture with needles pre-washed with 0.5M EDTA, after which the mouse was perfused with ice cold PBS prior to harvesting other tissues. To isolate inflammatory cells from spinal cords, individual spinal cords were dissociated through a wire mesh after which leukocytes were isolated using a Percoll (GE Healthcare Life Sciences) gradient, collecting leukocytes at the 37/90% Percoll interface. Cell suspensions were generated from lymphoid tissues by first dissociating them between frosted glass slides. The spleen and blood were then lysed for 2 min at 37°C to remove red blood cells with ACK buffer. Dead cells were identified by staining with a Fixable Viability Dye eFluor506 (eBioscience) according to manufacturer’s protocol. All cells were then blocked with an anti-Fcγ receptor, CD16/32 2.4G2 (BD biosciences), in PBS containing 2% FBS for 30 min on ice. Cells were then stained on ice for 30 mins with the listed combination of staining Abs, followed by a secondary stain with streptavidin for 15 mins on ice where necessary. Spleen cells were fixed in 2% PFA in PBS prior to running cells to prevent cell clumping. Flow cytometry was performed on a BD Immunocytometry Systems LSRII cytometer. Analysis was then completed using FlowJo software (TreeStar).

### Immunofluorescent histology

Spinal cord and lymph node tissue were prepared for histology as previously described (20). Briefly, at the end of the experiment or earlier, if mice reached a predetermined endpoint lymph node sections and spinal cord tissue spanning the cervical to lumbar regions were isolated and fixed in PLP. Spinal cords were then cut into five to nine evenly spaced sections and frozen in OCT (TissueTek) media. Tissue was then cut in serial cryostat sections at 7μm. Prior to staining, all slide-mounted tissue sections were blocked with PBS containing 1% BSA, 0.1% Tween-20, and 10% rat serum. After staining, sections were mounted with ProLong Gold Antifade Reagent (Invitrogen). When staining for MBP, tissue sections were additionally pre-treated to remove lipids in a series of ethanol gradients (0%, 50%, 70%, 90%, 95%, 100%, 95%, 90%, 70%, 50% and 0% ethanol in deionized water for one minute at each concentration) prior to the blocking step. Tiled images of whole spinal cord sections were collected using a DM5500B fluorescence microscope (Leica) at 20x.

### mMOG_tag_ immunization and cell isolation for transfer

To induce activated anti-MOG T and B cells for subsequent transfer into chronic sEAE mice, 2D2 FRP^+^ mice were immunized with a fusion protein antigen based on the extracellular domain of mouse MOG protein (mMOG_tag_) (18, 34) as previously described (33). Briefly, 6-8 week old 2D2 FRP^+^ mice were immunized with 0.5 mg of mMOG_tag_ in CFA (Sigma-Aldrich) s.c. at two sites near the base of the tail. 5d post immunization, draining inguinal lymph nodes were harvested and dissociated into transfer buffer (10 mM HEPES, 25 μg/mL gentamycin, 2.5% Acid citrate dextrose solution A in PBS). Suspended and filtered cells were injected i.v. into the tail vein of chronic sEAE mice at a 1:1 ratio of donor to recipient mice.

### Imaging and Statistical analysis

Microscopy images were analyzed using Fiji software. The number of infiltrating B and T cells were determined by cell counting the number of cells per spinal cord area. The total area of the spinal cord was determined by either FluoroMyelin staining or glial fibrillary acidic protein (GFAP) staining, while regions of demyelination were determined by the absence of MBP. The percent of demyelination was calculated by dividing the area of demyelination to the total area of the spinal cord. To determine the size of the cluster area, I looked at regions of the meninges (determined by the lack of GFAP staining) that contained B and T cell infiltration. PRISM software (GraphPad, La Jolla, California) was used for all statistical analysis. A Student t test was used for single comparisons, and ANOVA followed by a Student t test with Bonferroni correction was used for multiple comparisons.

## Results

### CNS-infiltrating B cells are reduced 7d post i.v. treatment with anti-mCD20

To determine the optimal dose of anti-mCD20 (5D2) to effectively deplete B cells in mice, we administered either 150μg or 250μg (based on previously published studies (35)) of anti-mCD20 (5D2) into wild type mice *i.v.* B cell numbers were analyzed in the blood, peripheral lymph nodes and spleen by flow cytometry 1d post treatment. As expected, B cells were almost entirely undetectable in the blood (**Figure 1A**), demonstrating that depletion of B cells in the circulation is rapid and effective. B cell numbers in lymph nodes (axial, brachial and inguinal), and the spleen were also reduced, but not to the same degree as in the blood. This suggests that anti-mCD20 that is administered *i.v.* has limited access to B cells in tissues compared to cells in the circulation, consistent with previous analysis of *i.p.* administered anti-mCD20 (26). As no difference was observed between the two doses, all subsequent experiments used 150μg as the treatment dose.

**Figure 1.**
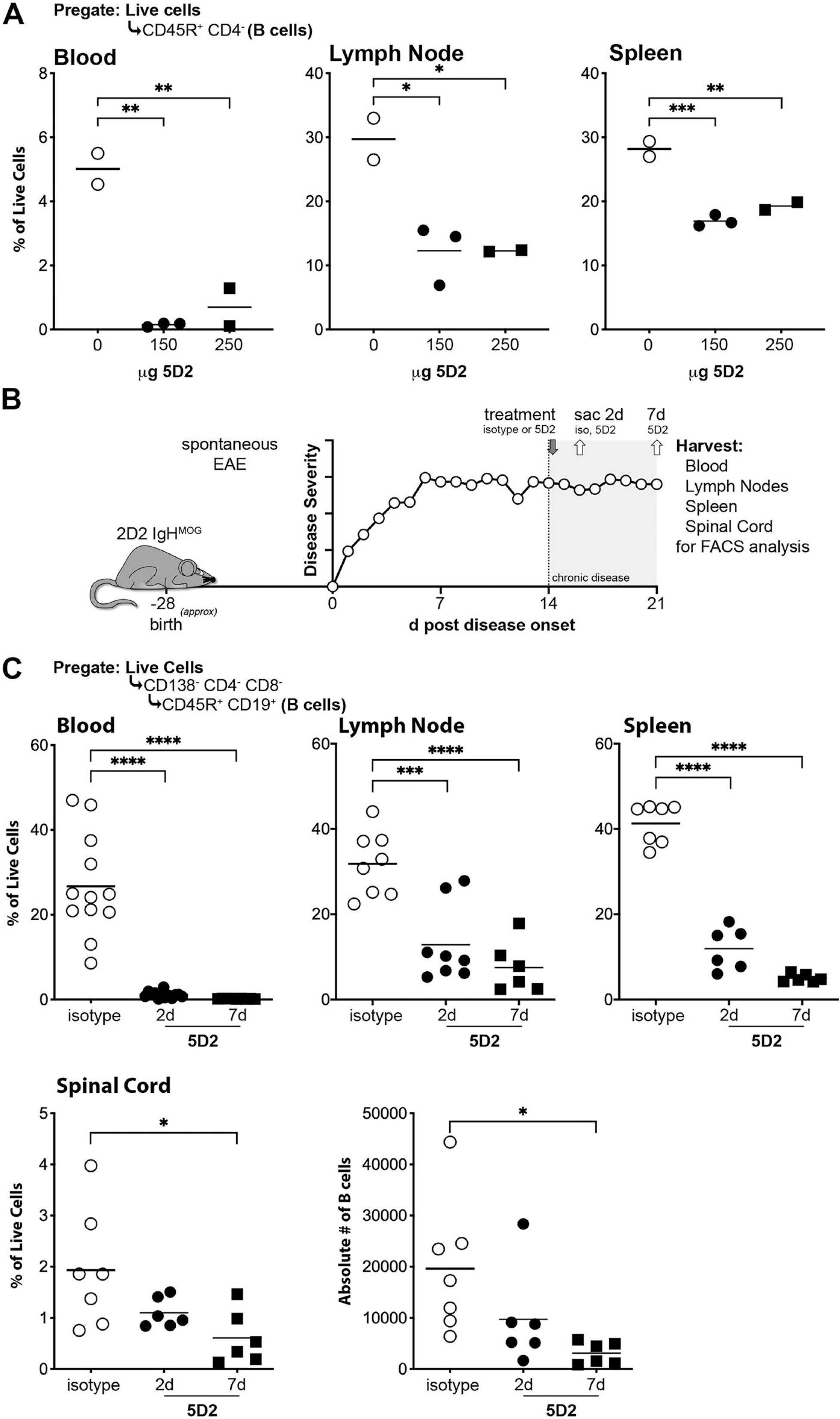
Depletion of meningeal B cells by systemically administered anti-mCD20 is delayed compared to other tissues. (**A**) Wild type mice were administered saline or either 150μg or 250μg anti-mCD20 (5D2) *i.v.* 1d post injection, blood, lymph nodes (axial, brachial and inguinal), and the spleen were harvested for flow cytometry analysis of B cell numbers. (**B**) 2D2 IgH^MOG^ sEAE mice in the chronic phase disease were treated with anti-mCD20 antibodies or isotype control. Tissue was harvested 2d or 7d later for analysis by flow cytometry. (**C**) Data is represented as percentage of live cells or absolute number of cells, as indicated. Each symbol represents and individual mouse. *p <0.05, **** p <0.0001.

To determine if systemic depletion of B cells also reduces the number of B cells in the inflamed spinal cord in sEAE, we administered anti-mCD20 *i.v.* during the chronic phase of disease (~2wks post onset, **Figure 1B**, see **Table 1** for the disease duration, severity, and sex of experimental mice). We have shown that T cell infiltration of the parenchyma and white matter demyelination is ongoing in this model at this timepoint, and also that B cells have established clusters in the meninges of the spinal cord (20). We analyzed B cell numbers in the blood, lymph nodes, spleen, and spinal cord by flow cytometry 2- or 7-days following treatment. As expected based on the above pilot study in healthy mice, B cells in the circulating blood were almost undetectable 2d post treatment and remained absent 7d post treatment, signifying that B cells were not replaced by hematopoiesis within this timeframe (**Figure 1C**). No decrease in CD4 or CD8 T cells were observed, as expected (**data not shown**). B cells were also significantly reduced in lymphoid tissues (lymph nodes and spleen) 2d and 7d post anti-mCD20-treatment compared to mice treated with isotype control antibody, but to a lesser extent than in the circulation (**Figure 1C**). In contrast, there was little indication of a reduction in B cells in the inflamed spinal cord by 2d pos treatment and it was not until 7d post treatment that a significant reduction in B cells was observed, both as a percentage of live cells and absolute number of B cells. This delay in the depletion of B cells in the inflamed CNS could be because it takes longer for the anti-mCD20 antibody to cross the blood brain barrier, or it could be because there is ongoing turnover of the CNS B cell population, and the elimination of circulating cells prevents the recruitment of replacement cells to the inflamed CNS.

**Table 1:** Phenotype of mice treated in flow cytometry experiments.

|  |  | **Average** | **STDEV** | **n** |
| --- | --- | --- | --- | --- |
| **Isotype**<br>**Control** | age of onset | 29.89 | 3.14 | 9 |
|  | d of onset | 15.11 | 5.95 |  |
|  | max score | 15.67 | 0.71 |  |
|  | end score | 14.33 | 1.66 |  |
|  | F (%) | 55.56 |  |  |
| **d2 5D2 Rx** | age of onset | 28.44 | 2.88 | 9 |
|  | d of onset | 14.44 | 5.90 |  |
|  | max score | 14.89 | 1.27 |  |
|  | end score | 12.67 | 2.24 |  |
|  | F (%) | 55.56 |  |  |
| **d7 5D2 Rx** | age of onset | 32.67 | 1.86 | 6 |
|  | d of onset | 17.83 | 5.95 |  |
|  | max score | 14.83 | 1.33 |  |
|  | end score | 11.67 | 3.20 |  |
|  | F (%) | 33.33 |  |  |

**Table 2:** Phenotype of mice treated in immunofluorescence experiments.

|  |  | Average | STDEV | n |
| --- | --- | --- | --- | --- |
| **Control** | **age of onset** | 29.25 | 3.105295 |  |
|  | **d of onset** | 17.5 | 4.14 |  |
|  | **max score** | 15.3 | 0.71 | 8 |
|  | **end score** | 13.8 | 1.75 |  |
|  | **F (%)** | 50 |  |  |
| **d7 5D2**<br>**Rx** | **age of onset** | 27.8 | 3.63 |  |
|  | **d of onset** | 22.2 | 1.30 |  |
|  | **max score** | 15.2 | 0.84 | 5 |
|  | **end score** | 10.8 | 2.28 |  |
|  | **F (%)** | 40 |  |  |

### Anti-mCD20 administration reduces the number of B cells in the meninges and this corresponds with less demyelination

Because meningeal B cell clusters correlate with disease severity in MS (14, 15) and EAE (19, 20), we performed additional experiments to determine if systemic B cell depletion reduces B cell accumulation in the meninges and also spinal cord pathology in chronic sEAE. sEAE mice in the chronic phase of disease were administered anti-mCD20 *i.v.* and spinal cords were harvested 7d later for histological analysis (see **Table 2** for disease parameters). Initial analysis revealed no differences in any measured parameters between isotype control treated mice and untreated mice, and therefore both were included in the control group.

As we observed previously (20), clusters of B cells were evident in the spinal cord meninges of chronic sEAE mice, and these clusters typically formed adjacent to regions of T cell infiltration into the spinal cord white matter and regions of demyelination (**Figure 2A**). Most clusters were a mixture of B cells and T cells, however in some circumstances the largest clusters showed evidence of organized separation of these cell types (as shown in **Figure 2A**). Examples of organized and disorganized clusters could be found in both control and anti-mCD20-treated mice. However, and consistent with the reduction in CNS B cells observed by flow cytometry (**Figure 1C**) above, treatment significantly reduced the density of B cells within meningeal clusters compared to untreated controls (**Figure 2B**). Absolute numbers of meningeal B cells were also lower, but this did not reach significance. Interestingly, the area of the clusters themselves did not change over this time period (**Figure 2C**). Importantly, the area of white matter demyelination was significantly reduced in treated mice (**Figure 2D, E**). This did not translate to a significant difference in disease score within this 7d timeframe (**Figure 2F**), consistent with recent observations by others (36) who similarly observed improved tissue pathology after a longer-duration treatment with anti-CD20 that did not translate into a measurable change in disease severity using standard scoring systems. Nevertheless, our observations suggest that meningeal B cells contribute to local tissue pathology.

**Figure 2:**
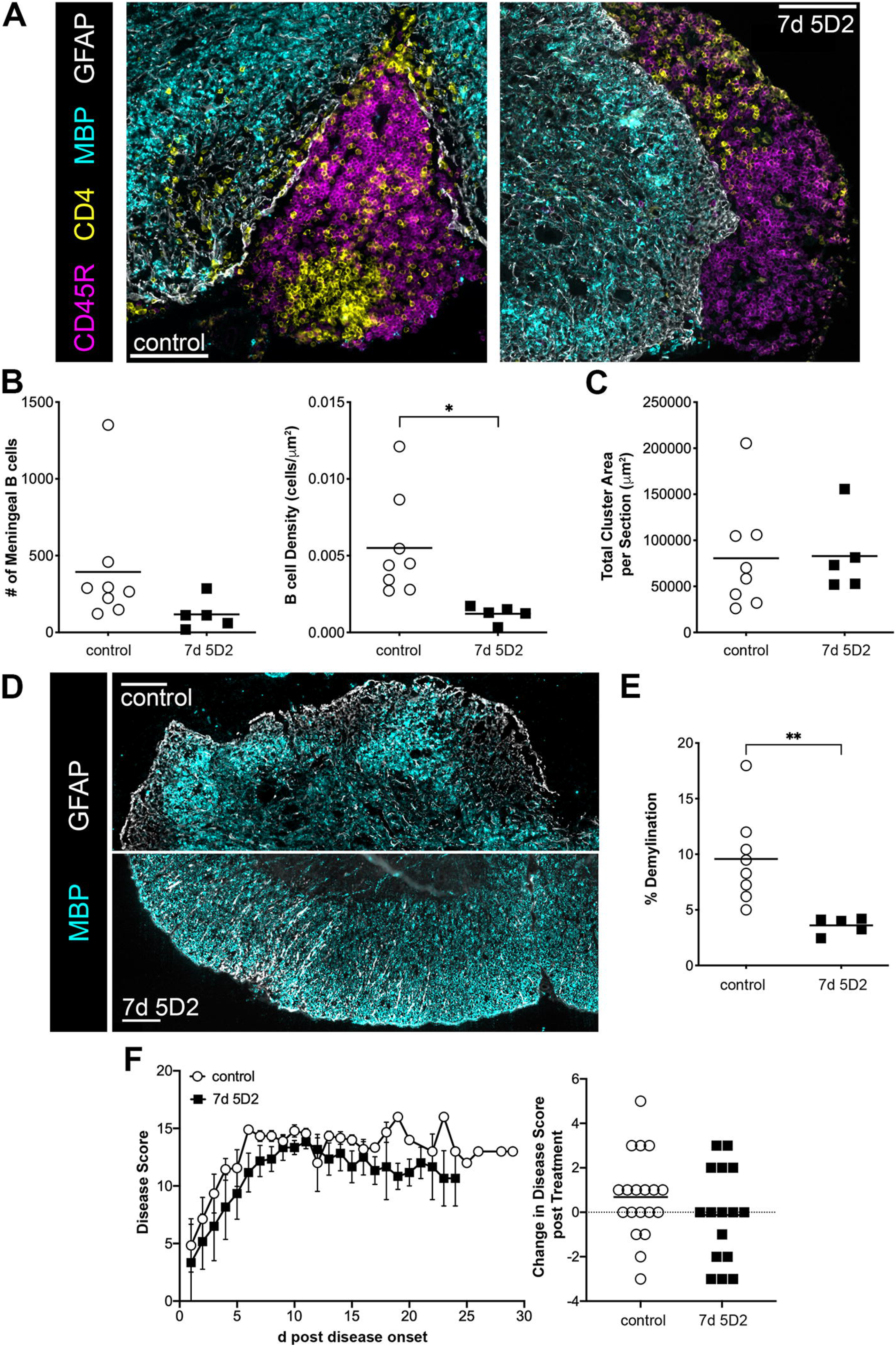
Anti-mCD20 treatment reduces meningeal B cell infiltration and demylination by 7d post treatment. (**A**) Representative immunofluorescence images of spinal cords from an isotype-control treated mouse (*left*) and d7 anti-mCD20 treated mouse (*right*). These examples were chosen to represent especially large B cell clusters, although the majority were smaller and showed less evidence of separation between B cells and T cells (see Figures 3 and 4 for additional examples). In the case of the control spinal cord (*left*), extensive T cell infiltration and demyelination of the adjacent white matter is apparent. (**B**) The number and density of B cells in the meninges, as well as total cluster area (**C**-defined by the presence of B and T cells), was determined from multiple sections from each mouse. (**D**) Representative images showing extensive demyelination in a control sEAE mouse (*top*) and less demyelination in a 5D2-treated sEAE mouse (*bottom*). (**E**) Area of demyelination was determined by measuring regions staining with GFAP but not MBP and expressed as a percentage of the total area of the spinal cord section. Each symbol represents an individual mouse. *p <0.05, **p <0.01. (**F**) Disease severity of control and anti-mCD20 treated mice used in the histological analysis above was tracked (****left****), along with the relative change in disease score over the final 7d prior to sacrifice (****right****). Data is shown as mean +/- SEM.

### Reductions in meningeal B cells are not likely the result of direct anti-mCD20-mediated depletion

In the above experiments we established that depletion of B cells in the spinal cord following systemic administration of anti-mCD20 is delayed relative to that observed in the circulation and lymphatic tissues (**Figure 1C**). To determine if *i.v.* administered anti-mCD20 has differential access to B cells in the inflamed spinal cord compared to B cells in the circulating blood and lymphatic tissue, we used flow cytometry to identify B cells bound by *i.v.* administered 5D2 anti-mCD20 2 or 7d post treatment. Of the very few detectable B220^+^ CD19^+^ B cells remaining in the circulation 2d post treatment, approximately half were bound by 5D2 (**Figure 3A**). This suggests that B cells bound by 5D2 are rapidly eliminated in the blood, perhaps as they rejoin the circulation from peripheral tissues. In lymphoid tissue (spleen and lymph nodes), where measurable numbers of B cells were still evident (**Figure 1C**), the large majority of B cells were bound by 5D2 by d2 and remained so at d7. This may indicate that anti-mCD20-mediated depletion within tissues is slower than it is in the blood. In contrast, almost none of the B cells in the spinal cord were bound by 5D2 2d after *i.v.* administration, and this only increased to ~10% of CNS B cells 7d post administration. This suggests that 5D2 may not be able to cross the blood brain barrier to label cells already in the tissue, or that access to the CNS is much slower than other tissues.

**Figure 3:**
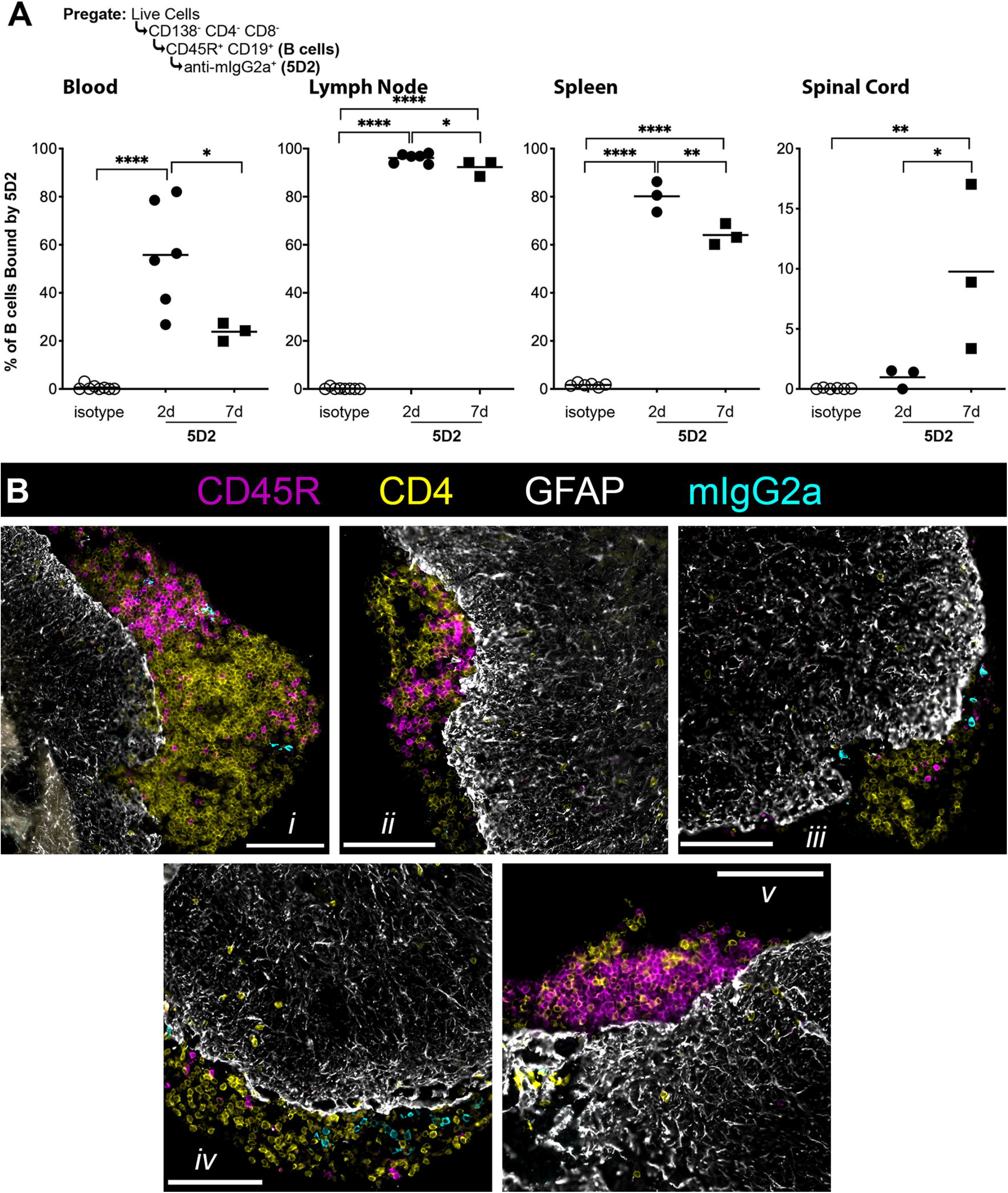
Meningeal B cell depletion is indirect and is likely the result of eliminating circulating cells. 2D2 IgH^MOG^ sEAE mice were administered isotype antibody or anti-mCD20 (5D2). (**A**) Circulating blood, lymph nodes (axial, brachial and inguinal), spleen and spinal cord were harvested 2d or 7d post treatment for analysis by flow cytometry. Anti-IgG2a was used to identify cells bound by 5D2. Each symbol represents an individual mouse. *p <0.05, **p <0.01, **** p <0.0001. (**B**) Representative immunofluorescence images of spinal cord meningeal B cell clusters a single chronic sEAE mouse (of 3) d7 post treatment with anti-mCD20 showing different patterns of 5D2 binding of cluster B cells. Panels *i* and *ii* are different clusters from the same section from the cervical spinal cord, and panels *iv* and *v* are different clusters from the same section from the thoracic spinal cord.

In order to differentiate between these possibilities, we performed a separate experiment to look by histology for direct evidence of B cells bound by 5D2 in the meninges, and also to determine the pattern of binding within meningeal clusters. If systemically administered anti-mCD20 does cross the blood brain barrier on its own by 7d post administration to directly bind B cells in meningeal clusters, we would expect to see all or most B cells within a given cluster to be evenly bound by the 5D2 antibody. This is not what we observed, however. Instead, 5D2-bound B cells were most commonly observed in meningeal clusters as individual, brightly-stained cells distributed among unlabeled cells (**Figure 3B**, panels *i*, *iii*, and *iv*). This pattern is more consistent with B cells being heavily labelled in the circulation and then carrying the anti-mCD20 antibody with them as they are recruited to the cluster. Interestingly, accumulation of 5D2-bound B cells was highly variable within individual clusters, where some clusters did not appear to have any (**Figure 3B**, panels *ii* and *v*) and, in rarer instances, most B cells were bound by 5D2 (panel *iv*). This inconsistency suggests that B cell recruitment and turnover rates differ between different clusters. Indeed, different patterns could be observed between clusters from the same tissue section (compare panel *i* to *ii*, and *iv* to *v*), and the degree of 5D2 binding corresponded to the degree of B cell elimination from the cluster. This suggests that anti-CD20 treatment may effectively eliminate B cells in “active” clusters with ongoing recruitment. Regardless, these observations together suggest that B cells encounter and are bound by systemically administered anti-CD20 in the circulation and are then recruited to the inflamed CNS, rather than the antibody itself crossing the blood meningeal barrier and binding cells for depletion directly in the tissue.

### T cells accumulate in the meninges following anti-mCD20 depletion of B cells

As noted above, the area of the meningeal lymphoid clusters did not change following anti-mCD20 treatment (**Figure 2C**), despite the fact that the density of B cells within the clusters was reduced (**Figure 2B**). Further analysis revealed that, with the decrease in B cells, CD4 T cells increased in the meninges of treated mice, resulting in a significant increase in both the numbers and density of these cells within clusters compared to control treated mice (**Figure 4 A and B**). Overall, this resulted in a significant shift in the dominant cell type within clusters from B cells to T cells (**Figure 4C**). This increase in CD4^+^ T cells is not likely due to increased proliferation, as there was no difference in the percentage of Ki67^+^ T cells as measured by microscopy (**Figure 4D**) or flow cytometry (**data not shown**). Flow cytometry also failed to show a difference in expression of IL-10, IFNγ, or IL-17 by CNS T cells (**data not shown**), again suggesting that the reduction in meningeal B cells did not impact T cell activation by this timepoint. Further, there was no change in the number of CD4^+^ T cells in the spinal cord white matter by d7 post treatment (**Figure 4E**), indicating that T cells do not leave the parenchyma to populate the meninges.

**Figure 4:**
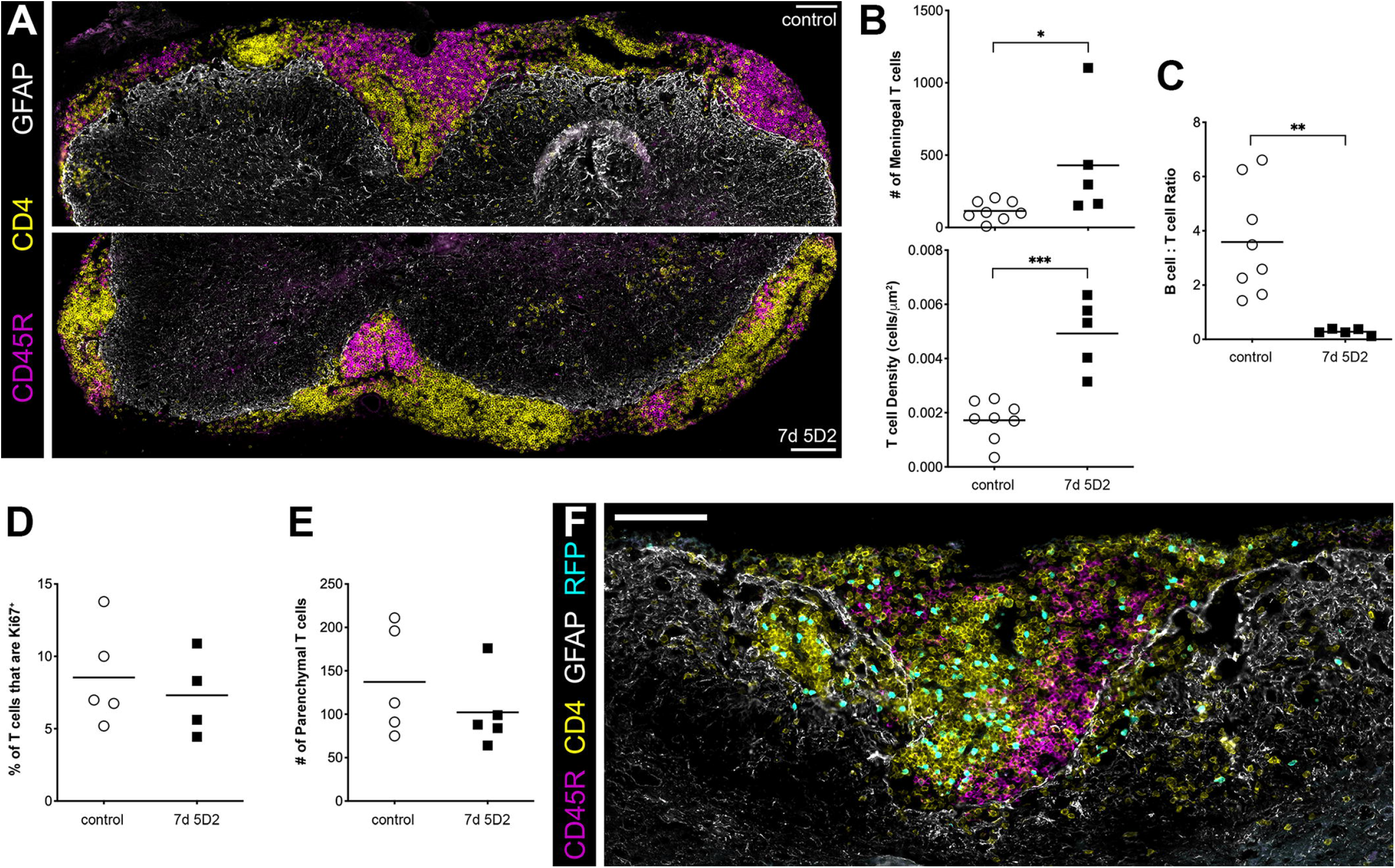
T cell infiltration of the meninges increases following B cell depletion. (**A**) A representative immunofluorescence image of a spinal cord from an isotype-control treated mouse (****top****) and a d7 anti-mCD20 (5D2) depleted mouse (****bottom****) showing a change in the dominant meningeal infiltrates from B cells to T cells. CD4^+^ cells and CD45R^+^ cells in the meninges were counted and the area of meningeal clusters was measured to determine the number and density of T cells in the clusters (**B**) and to calculate the ratio between B cells and T cells in the meninges (**C**). Separate sections were stained with Ki67 and CD4 to determine the percentage of CD4^+^ T cells in cell cycle (**D**). CD4^+^ T cells in the spinal cord parenchyma were counted (**E**). Each symbol represents an individual mouse. *p <0.05, **p <0.01. (**F**) sEAE mice in the chronic phase of disease were treated with anti-mCD20. 7d later, cells were isolated from the draining lymph nodes of RFP^+^ 2D2 mice that had been immunized with MOG protein 5d prior. These cells were transferred *i.v.* into the treated sEAE mice. 4d post transfer, spinal cords were harvested for histological imaging of transferred RFP^+^ cells into the inflamed spinal cords.

Finally, we transferred lymph node cells from MOG-immunized 2D2 RFP^+^ mice (5d post immunization) into chronic sEAE mice 7d post treatment with anti-mCD20. Spinal cords were harvested 4d post transfer to look for evidence of recruitment of the transferred RFP^+^ lymphocytes to the spinal cord. Interestingly, fewer RFP^+^ B cells were found in the lymph nodes of anti-mCD20-treated mice compared to untreated controls (flow cytometry data not shown), indicating that depletion of peripheral B cells was still effective 7-11d post treatment. RFP^+^ CD4^+^ cells, but not RFP^+^ CD45R^+^ B cells were readily apparent in meningeal clusters of anti-mCD20-treated mice by histology (**Figure 4F**), suggesting that extensive T cell recruitment to the inflamed CNS is ongoing following B cell depletion and that recruitment from the circulation is the likely source of the greater numbers of cluster T cells.

## Discussion

Here, we characterized the effects of anti-mCD20 depletion during the chronic phase of sEAE in 2D2 IgH^MOG^ mice. We found that B cell depletion in the inflamed spinal cord was delayed compared to peripheral tissues, including circulating blood, lymph nodes, and spleen. We attribute this delay to the mechanism of reduction of meningeal B cell numbers, which we believe results from the ongoing turnover of the population and the removal of the peripheral B cell pool from which it is replenished. Indeed, previous work from our lab shows that B cells are continuously recruited to meningeal clusters from the circulation at the chronic phase of EAE (33). Further, we did not find evidence that the 5D2 antibody crossed the blood brain barrier on its own, and our observations are more consistent with a scenario where B cells are bound by anti-CD20 in the periphery and carry the antibody with them as they are recruited into the inflamed CNS. Regardless, we consistently found that systemic administration of anti-mCD20 did ultimately result in fewer B cells in the CNS and specifically in the meninges of treated compared to control mice.

By focusing on the chronic phase of sEAE we show that systemically administered, anti-CD20 B cell-targeting therapies do reduce B cell numbers within already established lymphoid clusters in the inflamed CNS. Most previous studies investigating the effects of B cell depletion in models of CNS autoimmunity focused on the initiation and acute phase of disease. Treatment at the acute phase of disease or earlier would prevent the formation of meningeal clusters, and may also target other B cell populations with different roles in disease. Indeed, it is clear that depending on the model, B cell depletion before or shortly after disease induction results in less or more severe disease. For example, depleting B cells prior to disease onset in MOG_35-55_ peptide induced EAE results in increased disease severity (25, 26, 35) which is attributed to the loss of IL-10 production by B cell lineage cells (25). In contrast, when B cells are depleted prior to induction of EAE using larger protein antigens, disease severity is often reduced (26, 29, 30, 35, 37). Further, when B cells are depleted during the acute phase of recombinant MOG_1-117_ EAE (4d post EAE onset), frequencies of Th1, Th17 and regulatory T cells in the peripheral blood and CNS were decreased 14d post treatment (26), suggesting a role for B cells in modulating T cell responses in this early phase of disease, most likely from outside of the CNS.

Clusters of B cells in the meninges are associated with more severe disease in both humans (14, 15) and our sEAE model (20), and it follows that their reduction through anti-CD20 treatment coincided with a reduction in white matter demyelination in our studies. It is not clear if this is because demyelination was prevented from occurring in the first place, or if there was some opportunity for remyelination within the 7d treatment window. Remyelination following more extended anti-CD20 treatment in a different chronic model of EAE was observed by Breakell *et.al.* (36). The difference in white matter pathology following only 7d of treatment in our studies indicates that the therapeutic benefit of B cell depletion can be rapid, and that it is temporally associated with the reduction of B cells within established clusters in the meninges. Nevertheless, we cannot exclude the possibility that therapeutic benefit was through some other peripheral population. Still, unlike B cell depletion in the acute phase of disease discussed above, we did not observe any evidence of changes in cytokine production by T cells in our chronic model.

We were surprised to find that, while systemic administration of anti-CD20 did reduce the number of B cells in meningeal clusters, it also resulted in an increase in the number of T cells within the meninges. Typically, increased T cell infiltration of the CNS is associated with worse tissue pathology and disease. Further, we did not find evidence that the T cells in the meninges were less activated or more regulatory, based on levels of proliferation and cytokine production. Still, it must be emphasized that the increase in T cell numbers was limited to the meninges, as we observed no change in numbers of parenchymal, white matter T cells following anti-mCD20 treatment. Further, while we did observe a small number of the transferred T cells in white matter lesions of anti-mCD20-treated chronic sEAE mice, the vast majority remained confined to the meninges. We think that it is likely that, given time, the overall number of infiltrating T cells will go down, including those in the meninges. This would be in line with observations in human MS where B cell depletion is associated with less CNS inflammation over the months following treatment (1–5, 10, 11).

The identification of meningeal inflammatory foci in both secondary progressive (13–15) and primary progressive MS (38, 39) led to the hypothesis that the pathogenic autoimmune response is propagated from these locations. As stated, meningeal lesions have been correlated to accelerated disease severity (15), and increased axonal atrophy in the brain and spinal cord (16, 40), suggesting they are a deleterious site of inflammation. Here, we show in a mouse model that recapitulates these features of disease that these B cell clusters are susceptible to intervention through peripheral administration of anti-CD20. Further, we show that a reduction in these cells is associated with improved pathology, further supporting the hypothesis that they contribute pathologic mechanisms to local inflammation. Future studies will be required to determine the mechanistic action(s) of meningeal B cells in chronic CNS pathology.

## Acknowledgements

The authors would like to thank the veterinarians and animal care staff at the West Valley Barrier Facility for their excellent husbandry of our experimental animals. YT is the recipient of an endMS Doctoral Studentships from the Multiple Sclerosis Society of Canada (MSSOC). KAP is the recipient of an endMS Post-Doctoral Fellowship from MSSOC. This study was funded by operating grants from the Canadian Institutes of Health Research and the MSSOC.

## References

1. Bar-Or, A., P. Calabresi, D. Arnold, D. Arnlod, C. Markowitz, S. Shafer, L. Kasper, E. Waubant, S. Gazda, R. Fox, M. Panzara, N. Sarkar, S. Agarwal, and C. Smith. 2008. Rituximab in Relapsing-Remitting Multiple Sclerosis: A 72-Week, Open-Label, Phase I Trial. Ann. Neurol. 63: 395–400.

2. Hauser, S. L., D. L. Arnold, R. J. Fox, N. Sarkar, and C. H. Smith. 2008. B-Cell Depletion with Rituximab in Relapsing–Remitting Multiple Sclerosis. N Engl J Med 13.

3. Hauser, S. L., A. Bar-Or, G. Comi, G. Giovannoni, H.-P. Hartung, B. Hemmer, F. Lublin, X. Montalban, K. W. Rammohan, K. Selmaj, A. Traboulsee, J. S. Wolinsky, D. L. Arnold, G. Klingelschmitt, D. Masterman, P. Fontoura, S. Belachew, P. Chin, N. Mairon, H. Garren, and L. Kappos. 2017. Ocrelizumab versus Interferon Beta-1a in Relapsing Multiple Sclerosis. N. Engl. J. Med. 376: 221–234.

4. Montalban, X., S. L. Hauser, L. Kappos, D. L. Arnold, A. Bar-Or, G. Comi, J. de Seze, G. Giovannoni, H.-P. Hartung, B. Hemmer, F. Lublin, K. W. Rammohan, K. Selmaj, A. Traboulsee, A. Sauter, D. Masterman, P. Fontoura, S. Belachew, H. Garren, N. Mairon, P. Chin, and J. S. Wolinsky. 2017. Ocrelizumab versus Placebo in Primary Progressive Multiple Sclerosis. N. Engl. J. Med. 376: 209–220.

5. Wolinsky, J. S., D. L. Arnold, B. Brochet, H.-P. Hartung, X. Montalban, R. T. Naismith, M. Manfrini, J. Overell, H. Koendgen, A. Sauter, I. Bennett, S. Hubeaux, L. Kappos, and S. L. Hauser. 2020. Long-term follow-up from the ORATORIO trial of ocrelizumab for primary progressive multiple sclerosis: a post-hoc analysis from the ongoing open-label extension of the randomised, placebo-controlled, phase 3 trial. Lancet Neurol. S1474442220303422.

6. Monson, N. L., P. D. Cravens, E. M. Frohman, K. Hawker, and M. K. Racke. 2005. Effect of Rituximab on the Peripheral Blood and Cerebrospinal Fluid B Cells in Patients With Primary Progressive Multiple Sclerosis. Arch. Neurol. 62: 258.

7. Stüve, O., S. Cepok, B. Elias, A. Saleh, H.-P. Hartung, B. Hemmer, and B. C. Kieseier. 2005. Clinical Stabilization and Effective B-Lymphocyte Depletion in the Cerebrospinal Fluid and Peripheral Blood of a Patient With Fulminant Relapsing-Remitting Multiple Sclerosis. Arch. Neurol. 62.

8. Piccio, L., R. T. Naismith, K. Trinkaus, R. S. Klein, B. J. Parks, J. A. Lyons, and A. H. Cross. 2010. Changes in B- and T-Lymphocyte and Chemokine Levels With Rituximab Treatment in Multiple Sclerosis. Arch. Neurol. 67.

9. Cross, A. H., J. L. Stark, J. Lauber, M. J. Ramsbottom, and J.-A. Lyons. 2006. Rituximab reduces B cells and T cells in cerebrospinal fluid of multiple sclerosis patients. J. Neuroimmunol. 180: 63–70.

10. Kappos, L., D. Li, P. A. Calabresi, P. O’Connor, A. Bar-Or, F. Barkhof, M. Yin, D. Leppert, R. Glanzman, J. Tinbergen, and S. L. Hauser. 2011. Ocrelizumab in relapsing-remitting multiple sclerosis: a phase 2, randomised, placebo-controlled, multicentre trial. The Lancet 378: 1779–1787.

11. Alping, P., T. Frisell, L. Novakova, P. Islam-Jakobsson, J. Salzer, A. Björck, M. Axelsson, C. Malmeström, K. Fink, J. Lycke, A. Svenningsson, and F. Piehl. 2016. Rituximab versus fingolimod after natalizumab in multiple sclerosis patients: Rituximab vs Fingolimod. Ann. Neurol. 79: 950–958.

12. Hauser, S. L., A. Bar-Or, J. A. Cohen, G. Comi, J. Correale, P. K. Coyle, A. H. Cross, J. de Seze, D. Leppert, X. Montalban, K. Selmaj, H. Wiendl, C. Kerloeguen, R. Willi, B. Li, A. Kakarieka, D. Tomic, A. Goodyear, R. Pingili, D. A. Häring, K. Ramanathan, M. Merschhemke, and L. Kappos. 2020. Ofatumumab versus Teriflunomide in Multiple Sclerosis. N. Engl. J. Med. 383: 546–557.

13. Serafini, B., B. Rosicarelli, R. Magliozzi, E. Stigliano, and F. Aloisi. 2004. Detection of Ectopic B-cell Follicles with Germinal Centers in the Meninges of Patients with Secondary Progressive Multiple Sclerosis. Brain Pathol. 14: 164–174.

14. Magliozzi, R., O. Howell, A. Vora, B. Serafini, R. Nicholas, M. Puopolo, R. Reynolds, and F. Aloisi. 2006. Meningeal B-cell follicles in secondary progressive multiple sclerosis associate with early onset of disease and severe cortical pathology. Brain 130: 1089–1104.

15. Howell, O. W., C. A. Reeves, R. Nicholas, D. Carassiti, B. Radotra, S. M. Gentleman, B. Serafini, F. Aloisi, F. Roncaroli, R. Magliozzi, and R. Reynolds. 2011. Meningeal inflammation is widespread and linked to cortical pathology in multiple sclerosis. Brain 134: 2755–2771.

16. Reali, C., R. Magliozzi, F. Roncaroli, R. Nicholas, O. W. Howell, and R. Reynolds. 2020. B cell rich meningeal inflammation associates with increased spinal cord pathology in multiple sclerosis. Brain Pathol. 30: 779–793.

17. Peters, A., L. A. Pitcher, J. M. Sullivan, M. Mitsdoerffer, S. E. Acton, B. Franz, K. Wucherpfennig, S. Turley, M. C. Carroll, E. Bettelli, and V. K. Kuchroo. 2011. Th17 Cells Induce Ectopic Lymphoid Follicles in Central Nervous System Tissue Inflammation. 20.

18. Dang, A. K., R. W. Jain, H. C. Craig, and S. M. Kerfoot. 2015. B cell recognition of myelin oligodendrocyte glycoprotein autoantigen depends on immunization with protein rather than short peptide, while B cell invasion of the CNS in autoimmunity does not. J. Neuroimmunol. 278: 73–84.

19. Kuerten, S., A. Schickel, C. Kerkloh, M. S. Recks, K. Addicks, N. H. Ruddle, and P. V. Lehmann. 2012. Tertiary lymphoid organ development coincides with determinant spreading of the myelin-specific T cell response. Acta Neuropathol. (Berl.) 124: 861–873.

20. Dang, A. K., Y. Tesfagiorgis, R. W. Jain, H. C. Craig, and S. M. Kerfoot. 2015. Meningeal Infiltration of the Spinal Cord by Non-Classically Activated B Cells is Associated with Chronic Disease Course in a Spontaneous B Cell-Dependent Model of CNS Autoimmune Disease. Front. Immunol. 6.

21. Batoulis, H., M. Wunsch, J. Birkenheier, A. Rottlaender, V. Gorboulev, and S. Kuerten. 2015. Central nervous system infiltrates are characterized by features of ongoing B cell-related immune activity in MP4-induced experimental autoimmune encephalomyelitis. Clin. Immunol. 158: 47–58.

22. Serafini, B., M. Severa, S. Columba-Cabezas, B. Rosicarelli, C. Veroni, G. Chiappetta, R. Magliozzi, R. Reynolds, E. M. Coccia, and F. Aloisi. 2010. Epstein-Barr Virus Latent Infection and BAFF Expression in B Cells in the Multiple Sclerosis Brain: Implications for Viral Persistence and Intrathecal B-Cell Activation. J. Neuropathol. Exp. Neurol. 69: 677–693.

23. Aloisi, F., and R. Pujol-Borrell. 2006. Lymphoid neogenesis in chronic inflammatory diseases. Nat. Rev. Immunol. 6: 205–217.

24. Haugen, M., J. L. Frederiksen, and M. Degn. 2014. B cell follicle-like structures in multiple sclerosis-with focus on the role of B cell activating factor. J. Neuroimmunol. 273: 1–7.

25. Matsushita, T., K. Yanaba, J.-D. Bouaziz, M. Fujimoto, and T. F. Tedder. 2008. Regulatory B cells inhibit EAE initiation in mice while other B cells promote disease progression. J. Clin. Invest. JCI36030.

26. Weber, M. S., T. Prod’homme, J. C. Patarroyo, N. Molnarfi, T. Karnezis, K. Lehmann-Horn, D. M. Danilenko, J. Eastham-Anderson, A. J. Slavin, C. Linington, C. C. A. Bernard, F. Martin, and S. S. Zamvil. 2010. B-cell activation influences T-cell polarization and outcome of anti-CD20 B-cell depletion in central nervous system autoimmunity: B Cells in CNS Autoimmunity. Ann. Neurol. 68: 369–383.

27. Bettelli, E., D. Baeten, A. Jäger, R. A. Sobel, and V. K. Kuchroo. 2006. Myelin oligodendrocyte glycoprotein–specific T and B cells cooperate to induce a Devic-like disease in mice. J. Clin. Invest. 116: 2393–2402.

28. Krishnamoorthy, G., H. Lassmann, H. Wekerle, and A. Holz. 2006. Spontaneous opticospinal encephalomyelitis in a double-transgenic mouse model of autoimmune T cell/B cell cooperation. J. Clin. Invest. 116: 2385–2392.

29. Monson, N. L., P. Cravens, R. Hussain, C. T. Harp, M. Cummings, M. de Pilar Martin, L.-H. Ben, J. Do, J.-A. Lyons, A. Lovette-Racke, A. H. Cross, M. K. Racke, O. Stüve, M. Shlomchik, and T. N. Eagar. 2011. Rituximab Therapy Reduces Organ-Specific T Cell Responses and Ameliorates Experimental Autoimmune Encephalomyelitis. PLoS ONE 6: e17103.

30. Pol, S., S. Liang, F. Schweser, R. Dhanraj, A. Schubart, M. Preda, M. Sveinsson, D. P. Ramasamy, M. G. Dwyer, G. Weckbecker, and R. Zivadinov. 2020. Subcutaneous anti-CD20 antibody treatment delays gray matter atrophy in human myelin oligodendrocyte glycoprotein-induced EAE mice. Exp. Neurol. 113488.

31. Lyons, J.-A., M. J. Ramsbottom, and A. H. Cross. 2002. Critical role of antigen-specific antibody in experimental autoimmune encephalomyelitis induced by recombinant myelin oligodendrocyte glycoprotein. Eur. J. Immunol. 32: 1905–1913.

32. Jain, R. W., K. A. Parham, Y. Tesfagiorgis, H. C. Craig, E. Romanchik, and S. M. Kerfoot. 2018. Autoreactive, Low-Affinity T Cells Preferentially Drive Differentiation of Short-Lived Memory B Cells at the Expense of Germinal Center Maintenance. Cell Rep. 25: 3342–3355.e5.

33. Tesfagiorgis, Y., S. L. Zhu, R. Jain, and S. M. Kerfoot. 2017. Activated B Cells Participating in the Anti-Myelin Response Are Excluded from the Inflamed Central Nervous System in a Model of Autoimmunity that Allows for B Cell Recognition of Autoantigen. J. Immunol. 199: 449–457.

34. Jain, R. W., A. K. Dang, and S. M. Kerfoot. 2016. Simple and Efficient Production and Purification of Mouse Myelin Oligodendrocyte Glycoprotein for Experimental Autoimmune Encephalomyelitis Studies. J. Vis. Exp. JoVE e54727–e54727.

35. Häusler, D., S. Häusser-Kinzel, L. Feldmann, S. Torke, G. Lepennetier, C. C. A. Bernard, S. S. Zamvil, W. Brück, K. Lehmann-Horn, and M. S. Weber. 2018. Functional characterization of reappearing B cells after anti-CD20 treatment of CNS autoimmune disease. Proc. Natl. Acad. Sci. 115: 9773–9778.

36. Breakell, T., S. Tacke, V. Schropp, H. Zetterberg, K. Blennow, E. Urich, and S. Kuerten. 2020. Obinutuzumab-Induced B Cell Depletion Reduces Spinal Cord Pathology in a CD20 Double Transgenic Mouse Model of Multiple Sclerosis. Int. J. Mol. Sci. 21: 6864.

37. Chen, D., S. J. Ireland, L. S. Davis, X. Kong, A. M. Stowe, Y. Wang, W. I. White, R. Herbst, and N. L. Monson. 2016. Autoreactive CD19+ CD20− Plasma Cells Contribute to Disease Severity of Experimental Autoimmune Encephalomyelitis. J. Immunol. 196: 1541–1549.

38. Choi, S. R., O. W. Howell, D. Carassiti, R. Magliozzi, D. Gveric, P. A. Muraro, R. Nicholas, F. Roncaroli, and R. Reynolds. 2012. Meningeal inflammation plays a role in the pathology of primary progressive multiple sclerosis. Brain 135: 2925–2937.

39. Bell, L., A. Lenhart, A. Rosenwald, C. M. Monoranu, and F. Berberich-Siebelt. 2020. Lymphoid Aggregates in the CNS of Progressive Multiple Sclerosis Patients Lack Regulatory T Cells. Front. Immunol. 10: 3090.

40. Magliozzi, R., O. W. Howell, C. Reeves, F. Roncaroli, R. Nicholas, B. Serafini, F. Aloisi, and R. Reynolds. 2010. A Gradient of neuronal loss and meningeal inflammation in multiple sclerosis. Ann. Neurol. 68: 477–493.

